# Constitutive UPR^ER^ activation sustains tumor cell differentiation

**DOI:** 10.1101/594630

**Authors:** Dimitrios Doultsinos, Mari McMahon, Konstantinos Voutetakis, Joanna Obacz, Raphael Pineau, Pierre-Jean Le Reste, Akram Obiedat, Juhi Samal, John B. Patterson, Qingping Zheng, Afshin Samali, Abhay Pandit, Boaz Tirosh, Aristotelis Chatziioannou, Eric Chevet, Tony Avril

**Author notes:** equal contribution (Mari McMahon, Konstantinos Voutetakis, Eric Chevet, Tony Avril). equal contribution and corresponding authors TA and EC.

## Abstract

Endoplasmic Reticulum (ER) proteostasis control and the Unfolded Protein Response (UPR^ER^) have been shown to contribute to tumor development and aggressiveness. As such, the UPR^ER^ sensor IRE1α (referred to as IRE1 hereafter) is a major regulator of glioblastoma (GBM) development and is an appealing therapeutic target. To document IRE1 suitability as an antineoplastic pharmacological target, we investigated how this protein contributed to GBM cell reprogramming, a property involved in treatment resistance and disease recurrence. Probing the IRE1 activity molecular signature on transcriptome datasets of human tumors, showed that high IRE1 activity correlated with low expression of the main GBM stemness transcription factors SOX2, SALL2, POU3F2 and OLIG2. Henceforth, this phenotype was pharmacologically and genetically recapitulated in immortalized and primary GBM cell lines as well as in mouse models. We demonstrated that constitutive activation of the IRE1/XBP1/miR148a signaling axis repressed the expression of SOX2 and led to maintenance of a differentiation phenotype in GBM cells. Our results describe a novel role for IRE1 signaling in maintaining differentiated tumor cell state and highlight opportunities of informed IRE1 modulation utility in GBM therapy.

## Introduction

Endoplasmic Reticulum (ER) proteostasis is controlled by several molecular machines including ER-bound mRNA translation, translocation of newly synthesized proteins, protein folding and quality control, protein export or degradation (ERAD) (1, 2). These molecular machines are coordinated by the Unfolded Protein Response (UPR^ER^), an adaptive pathway aiming at restoring ER homeostasis (2, 3). The UPR^ER^ is transduced by three ER-resident sensors, PERK, ATF6 and IRE1. The latter is a serine/threonine kinase and endoribonuclease, which upon ER stress oligomerizes and trans-auto phosphorylates, mediating downstream signaling events that include JNK activation, the unconventional splicing of XBP1 mRNA to produce the transcription factor XBP1s and the degradation of a specific subset of miRNAs and mRNAs; a process called RNA regulated IRE1-dependent decay or RIDD (1). IRE1 was shown to be involved in cancer development in many instances (1–6) and to represent an appealing therapeutic target in triple negative breast cancers (5, 6). We demonstrated that IRE1 signaling controls glioblastoma (GBM) development and aggressiveness through both XBP1 mRNA splicing and RIDD (4). Indeed GBM cells are often subjected to high metabolic demand, hypoxic stress, accelerated cell cycle events and partially overcome these stresses through IRE1 signaling (2). As such IRE1 confers tumor cells with aggressive characteristics including i) the pro-tumoral remodeling of the tumor stroma with immune and endothelial cells and ii) high migration/invasion characteristics (4). Targeting the RNase activity of IRE1 negatively impacts tumor growth due to blocking of pro-survival cellular mechanisms mediated by XBP1s. Moreover, IRE1 is involved in invasion, growth and vascularization, having been shown to carry out dual and at times antagonistic functions through XBP1s and RIDD signaling respectively, in GBM development (7, 8). Moreover, GBM recurrence and therapeutic resistance can be attributed to the appearance of cancer stem cell (CSC) and differentiated-to-stem cell reprogramming capabilities (9). Genetic characteristics delineating tumor stem-like cells and differentiated tumor cells have been identified with nestin a prominent player alongside SOX2 in the former (10); and GFAP, VIM and YKL40 in the later (11). Interestingly, we observed that nestin is overexpressed in GBM cells expressing a dominant negative form of IRE1 (12), thus suggesting a link between IRE1 signaling and expression of stemness features.

Since IRE1 and its target, the transcription factor XBP1s were shown to play major roles in cellular differentiation (13) and are involved in GBM development, we postulated that IRE1 activity may contribute to GBM cell stemness regulation and thus we investigated the impact of IRE1 inhibition on the differentiation status of GBM cells, provided that cancer cell reprogramming contributes to antineoplastic treatment resistance and disease recurrence through cancer stem cells.

## Results

### IRE1 activity is associated with the differentiated status of GBM specimens

The previously identified IRE1 activity signature of 38 genes (IRE1_38; (4, 8)) was confronted to the TCGA GBM cohort to stratify patients in two groups of high and low IRE1 activity (4). Since we observed nestin expression in tumors deriving from U87 cells deficient for IRE1 signaling (12), we then used the IRE1_38 signature to investigate the putative IRE1-dependent expression of markers of GBM differentiation or stemness. We found that stem cell markers were upregulated, whilst differentiation markers downregulated in tumors exhibiting low IRE1 activity as compared with those with high activity (Figures 1A**, S1A-C**). This was representative in the case of stem markers BMI1, CD133 and nestin and differentiation markers SMA, vimentin and YKL40 (Figure 1B). Subsequently we tested whether IRE1 activity could impact on the expression of transcription factors (TFs) involved in GBM reprogramming. Indeed, we observed that stemness associated TFs were markedly decreased in tumors with high IRE1 activity (Figure 1C), the four starkest of which were OLIG2, POU3F2, SALL2 and SOX2 (Figure 1D). To further document this phenomenon in cell lines, we utilized commonly available U251 and U87, as well as primary RADH87 and RADH85 (14) GBM lines genetically modified to invalidate IRE1 signaling (4, 8). We observed a marked upregulation of OLIG2, POU3F2, SALL2 and SOX2 in the IRE1 signaling deficient lines compared to parental or wild-type IRE1 overexpressing cells (Figures 1E**, S1F**). As such we show that in patient tumors, high IRE1 activity correlates with a low expression level of reprogramming factors, which is recapitulated in both commonly available and primary GBM cell lines.

**Figure 1.**
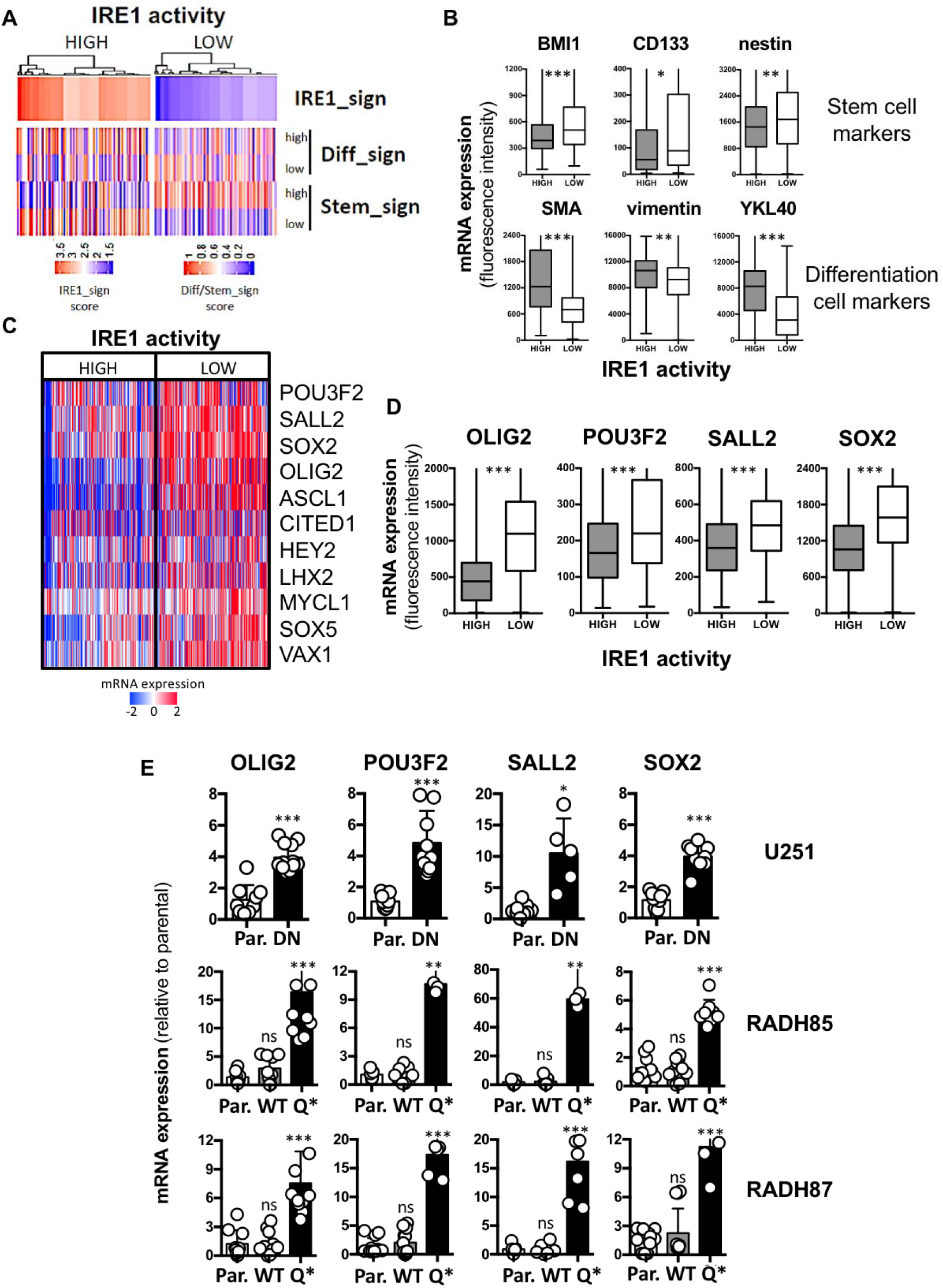
IRE1 activity is associated with cancer differentiated state in GBM specimens. **A)** Hierarchical clustering of GBM patients (TCGA cohort) based on high or low IRE1 activity confronted to differentiation and stem gene signatures derived from literature. **B)** mRNA expression of BMI1, CD133, nestin stem cell markers and SMA, vimentin, YKL40 differentiated cell markers based on microarray fluorescence intensity in high and low IRE1 activity tumors (TCGA cohort). **C)** Hierarchical clustering of GBM patients (TCGA cohort) based on high or low IRE1 activity confronted to a reprogramming TFs signature derived from literature. **D)** mRNA levels of reprogramming TFs OLIG2, POU3F2, SALL2, and SOX2 based on microarray fluorescence intensity in high and low IRE1 activity tumors (TCGA cohort). **E)** mRNA levels of reprogramming TFs in U251, RADH85 and RADH87 lines expressing WT, DN or Q* forms of IRE1 normalized to parental. (ns): not significant; (*): p<0.05; (**): p<0.01; (***): p<0.001.

### Genetic and pharmacological perturbation of IRE1 disturbs GBM differentiated phenotype

To further investigate the effects of IRE1 inhibition on GBM differentiated-to-stem reprogramming, we developed a culture system where cell lines normally grown in adherent 10% FCS-containing media (RADH primary and U87/U251 commonly available cell lines) were seeded in FCS-free neurosphere culture media (Figure 2A) and cell viability, cell number, phenotype as well as the ability of cells to form spheres from single cells (clonogenicity) were determined (Figures 2**, S2**). No major phenotypic difference was seen between the parental cells and ones overexpressing a WT IRE1, however IRE1 signaling deficient cells readily formed spheres pertaining to a stem phenotype over several passages (Figures 2B-C**, S2A-D**). Next, we evaluated the expression of OLIG2, POU3F2, SALL2 and SOX2 in cells grown in those conditions. Whereas no difference between parental and wild-type IRE1 overexpressing cells was observed, IRE1 signaling deficient cells showed a uniform increase of those markers (Figures 2D**, S2E**), supporting the hypothesis that IRE1 signaling is instrumental for differentiated phenotype maintenance. This was further documented at the protein level where the stemness markers SOX2, nestin and A2B5 were all upregulated whilst the differentiation marker NG2 was downregulated in IRE1 signaling deficient cells (Figures 2E**, S2F**). Moreover, IRE1 signaling deficient cells were more clonogenic than both parental or IRE1 WT overexpressing lines (Figures 2F**, S2G**), compounding the stem phenotype in the absence of functional IRE1. We have thus demonstrated that IRE1 genetic perturbation disturbs the differentiated phenotype in commonly used and primary GBM lines. To further document this observation we used the salicylaldehyde IRE1 ribonuclease inhibitor MKC8866 (hereafter called MKC) (15). Interestingly, MKC treatment phenocopied the effects IRE1 signaling invalidation on the capacity of cells (Figure 3**, S3**). When compared to the DMSO control, U251, RADH85, RADH87 cells treated with MKC formed spheres more readily (Figures 3A-C**, S3B-D**), displayed higher mRNA levels of stem and reprogramming markers and lower mRNA levels of differentiation markers (Figures 3C**, S3F**), results that were confirmed at the protein level (Figures 3D**, S3G**). The clonogenic ability of the cells was also significantly upregulated in the presence of MKC (Figures 3E**, S3H**). As such we show that both genetic and pharmacological perturbations of IRE1 lead to a loss of differentiated phenotype whilst at the same time push cells towards a sphere stem-like phenotype.

**Figure 2.**
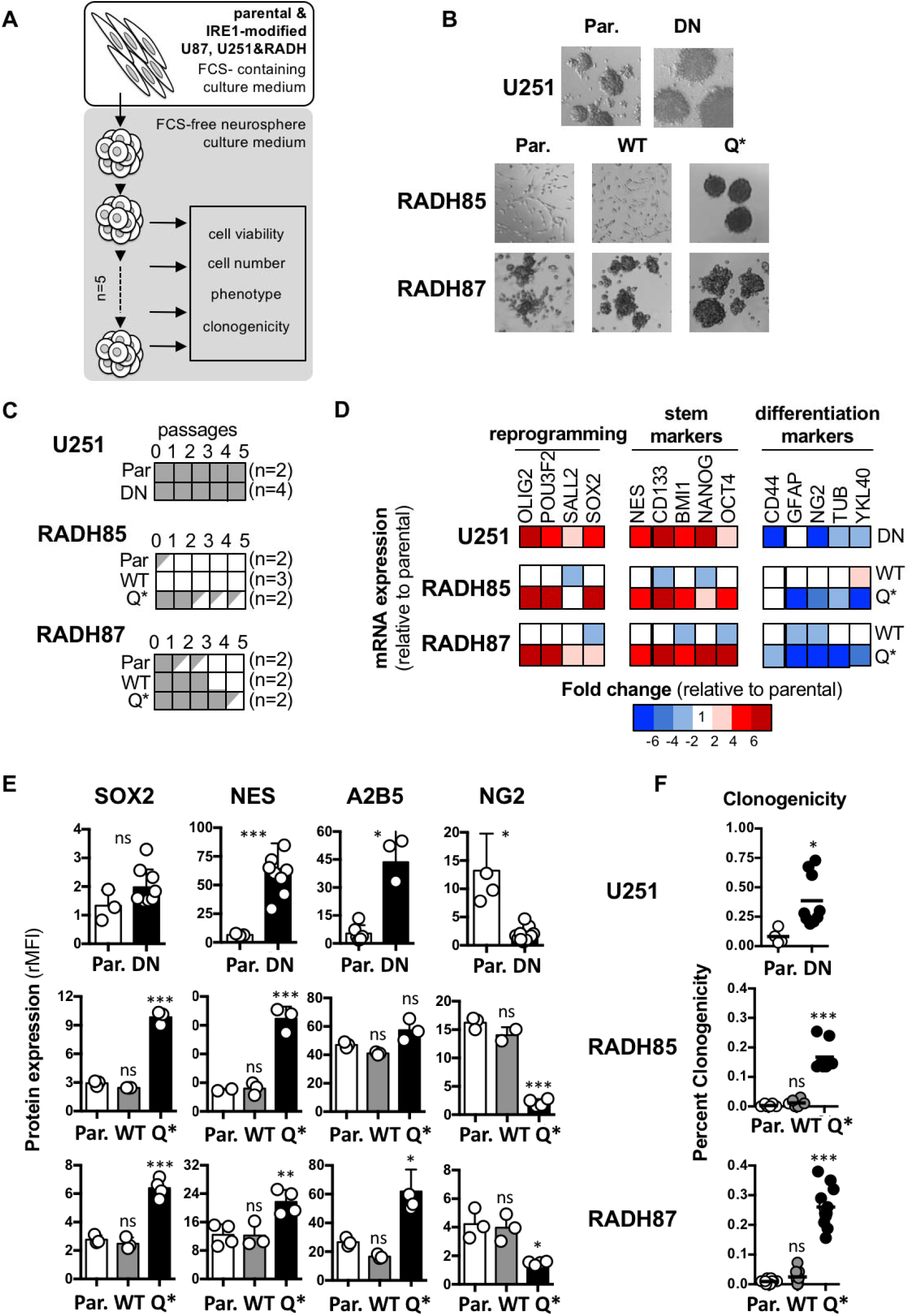
Genetic perturbation of IRE1 effect on GBM cell reprogramming. **A)** Schematic representation of GBM cell working model of differentiated to stem cell phenotype culture. **B)** Phenotypic characterization of U251/ RADH85/ RADH87 parental and overexpressing WT, DN or Q* forms of IRE1 when grown in neurosphere media. **C)** Differentiated GBM cell lines U251, RADH85 and RADH87 were cultured in neurosphere medium and were passaged every 14 days. If the number of cells was under the initial number of cells seeded (10^6^), the culture was stopped (n=2 to 4). **D)** Heat map representation of fold change of mRNA expression of genes involved in reprogramming, stemness and differentiation normalized to parental in U251, RADH85, RADH87 lines expressing WT, DN or Q* forms of IRE1 when grown in neurosphere media. **E)** Protein expression of reprogramming, stemness and differentiation markers in these lines compared to parental lines determined by flow cytometry. **F)** Clonogenicity of differentiated lines expressing WT, DN or Q* forms of IRE1 compared to parental lines when grown in neurosphere media. (ns): not significant; (*): p<0.05; (**): p<0.01; (***): p<0.001.

**Figure 3.**
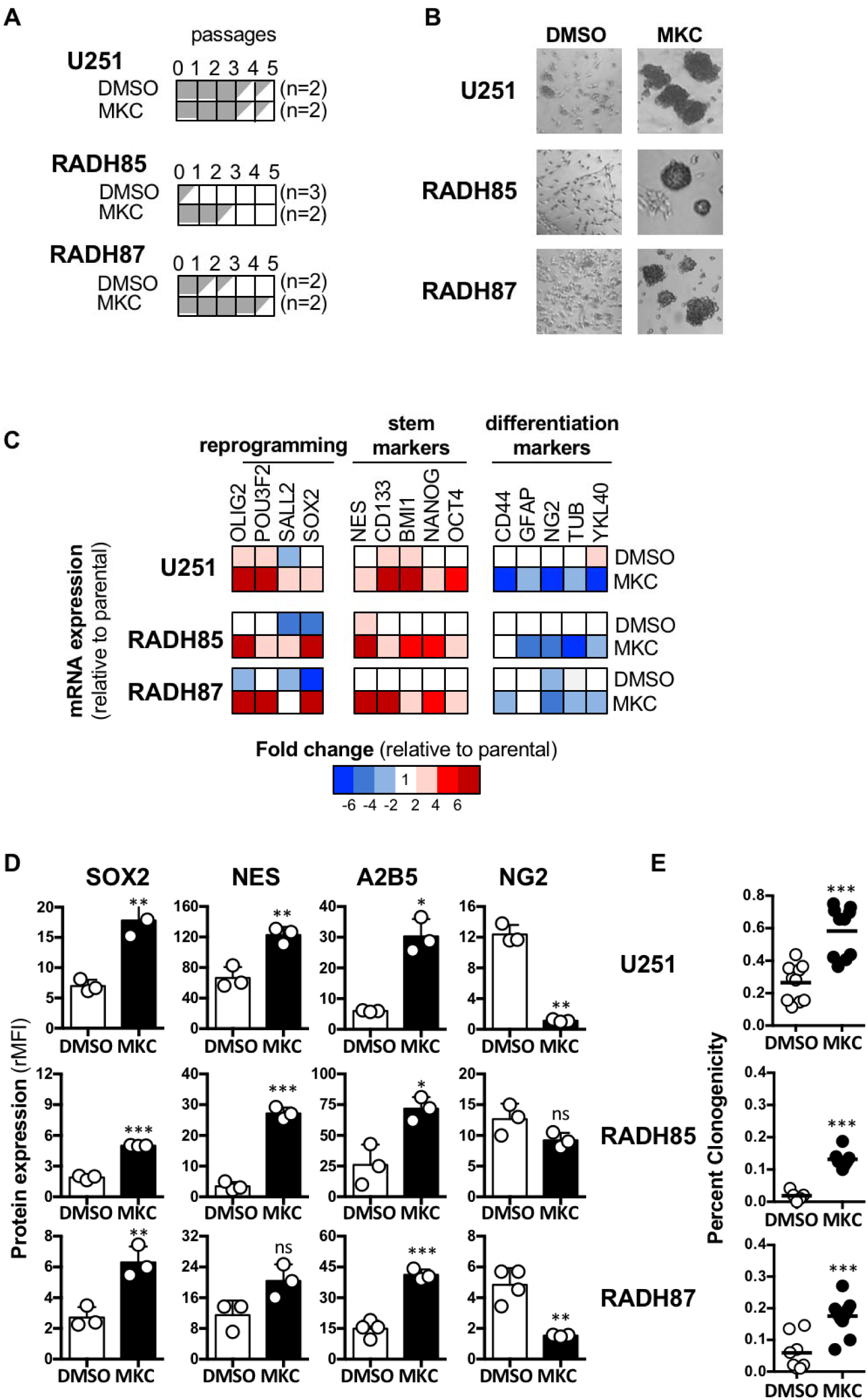
Pharmacological inhibition of IRE1 effect on GBM cell reprogramming. **A)** Differentiated GBM cell lines U251, RADH85 and RADH87 were cultured in neurosphere medium in the presence of MKC (5 µM), and were passaged every 14 days. If the number of cells was under the initial number of cells seeded (10^6^), the culture is stopped (n=2 to 3). **B)** Phenotypic characterization of parental adherent (RADH85/87 and U251) lines through culture in neurosphere medium treated with MKC or DMSO. **C)** Heat map representation of fold change of mRNA expression of genes involved in reprogramming, stemness and differentiation normalized to parental in U251, RADH85, and RADH87 lines when grown in neurosphere media in the presence of MKC or DMSO. **D)** Protein expression of reprogramming, stemness and differentiation markers in these lines compared to parental lines determined by flow cytometry. **E)** Quantification of clonogenicity of single cell parental, WT or Q* IRE1 expressing RADH85/87 and U251 lines when seeded in serum-free medium in the presence of MKC or DMSO. (ns): not significant; (*): p<0.05; (**): p<0.01; (***): p<0.001.

### IRE1 signaling modestly contribute to cancer stem cell differentiation

Since IRE1 activity appears instrumental for the maintenance of the differentiated phenotype in GBM cells, we next tested whether IRE1 plays a significant role in CSC differentiation. To this end, we utilized primary GBM lines selected in FCS-free neurosphere media (RNS lines; (16)) proficient or deficient for IRE1 signaling (4). These lines were seeded in 10% FCS-containing media in the presence of the bone morphogenetic factor BMP4 to induce differentiation (Figures 4A**, S4A**). Genetic perturbation of IRE1 signaling did not result in a significant phenotype change in RNS lines when grown in FCS-containing media in the presence of BMP4 (Figures 4B, **S4B-E**). Moreover, the differentiation markers GFAP, O4 and TUB showed slight decreases at the mRNA and protein levels in lines deficient for IRE1 signaling (Figure 4C and 4D). In the same line of observation, MKC treatment yielded no phenotypic changes in RNS cells in the presence of 10% FCS media and BMP4 (Figure 4E), whilst differentiation markers, although generally statistically significantly decreased at both the mRNA (Figure 4F) and protein level (Figure 4G), did not show the marked difference observed previously (Figures 2, 3). Hence, IRE1 signaling perturbation is modestly influencing cancer stem differentiation.

**Figure 4.**
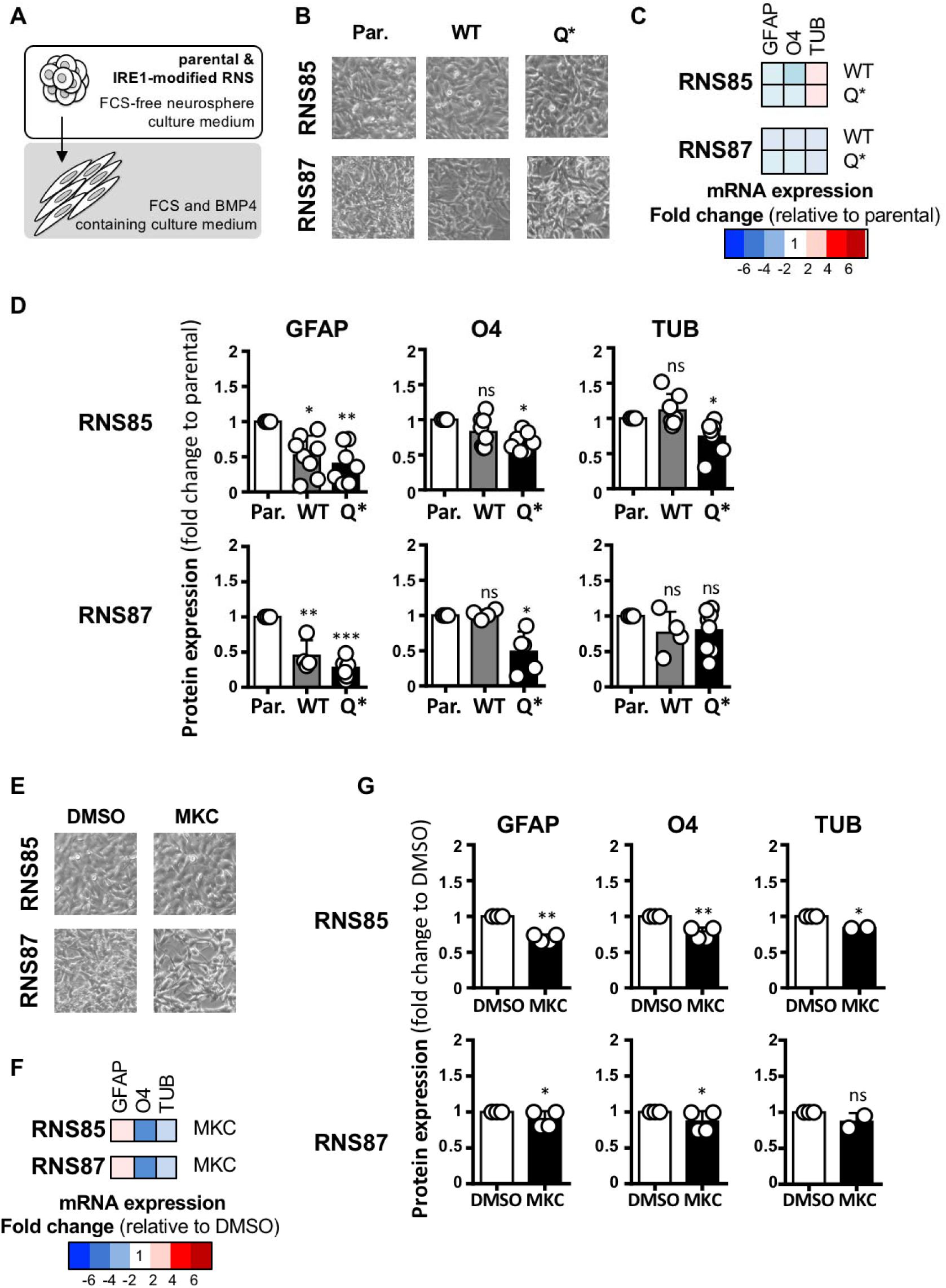
Role of IRE1 in GBM stem-to-differentiated state reprogramming. **A)** Schematic representation of GBM cell working model of stem-to-differentiated cell phenotype culture. **B)** Phenotypic characterization of RNS85/87 parental and overexpressing WT or Q* forms of IRE1 when grown in FCS and BMP4 containing media. **C)** Heat map representation of fold change of mRNA expression of genes involved in differentiation normalized to parental in RNS85, RNS87 lines expressing WT or Q* forms of IRE1 when grown in FCS and BMP4 containing media. **D)** Protein expression of differentiation markers in these lines normalized to parental determined by flow cytometry. **E)** Phenotypic characterization of parental RNS85/87 lines through culture in FCS and BMP4 containing medium treated with MKC or DMSO. **F)** Heat map representation of fold change of mRNA expression of genes involved in differentiation in parental RNS85, RNS87 lines when grown in FCS and BMP4 containing media in the presence of MKC or DMSO. **G)** Protein expression of differentiation markers in these lines in the presence of MKC or DMSO determined by flow cytometry. (ns): not significant; (*): p<0.05; (**): p<0.01; (***): p<0.001.

### Constitutive IRE1/XBP1s signaling maintains GBM cell differentiation

It is well described that IRE1 has a dual function mediated by its ribonuclease domain (3) through either XBP1 mRNA splicing or RIDD (17). We first correlated the expression of differentiation and stem markers in tumors stratified based on their XBP1s or RIDD status as previously defined (4). We found that high XBP1s activity tumors, expressed low levels of stem markers and high levels of differentiation markers (Figure 5A). This was confirmed by investigating individual genes as well. Stem markers BMI1, CD133 and nestin showed no significant difference between high XBP1s and high RIDD activity tumors whilst differentiated tumor cell markers SMA, vimentin and YKL40 were significantly more elevated in XBP1s high tumors as opposed to RIDD high tumors (Figure 5B). This led us to hypothesize that XBP1s rather than RIDD is the rate limiting factor in maintaining the differentiated GBM cell phenotype. Next, by monitoring the expression of 15 genes involved in reprogramming in XBP1s high or RIDD high tumors, we were able to discern that reprogramming factors were downregulated in TCGA tumors displaying high XBP1s activity (Figure 5C). This was statistically confirmed when measuring the expression of single genes OLIG2, POU3F2, SALL2 and SOX2 (Figure 5D). We then sought to evaluate whether XBP1s ablation was able to recapitulate this effect in commonly available U251 and primary (RADH85 and RADH87) adherent differentiated GBM cell lines. XBP1 silencing led to the significant upregulation of almost all 4 TFs across all human lines (Figures 5E, **S5**). We thus demonstrate that the IRE1-XBP1s signaling axis controls the ability of GBM cells to maintain differentiation.

**Figure 5.**
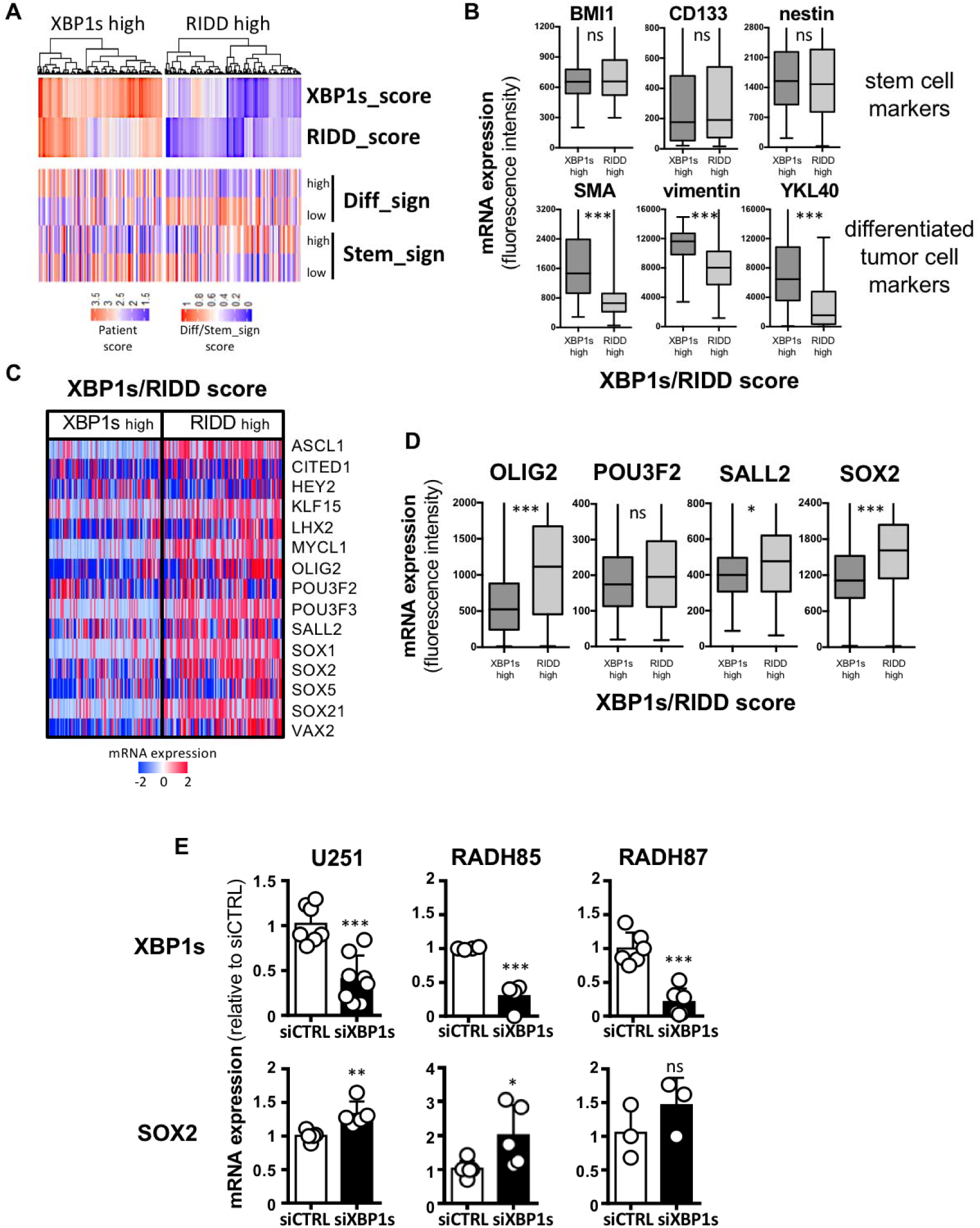
XBP1s involvement in GBM cell reprogramming. **A)** Hierarchical clustering of genes involved in differentiation and stemness in GBM (TCGA cohort) based on high XBP1s or high RIDD activity (blue low levels, red high levels). **B)** mRNA expression of BMI1, CD133, nestin stem cell markers and SMA, vimentin, YKL40 differentiated cell markers based on microarray fluorescence intensity in high XBP1s and high RIDD activity tumors (TCGA cohort). **C)** Hierarchical clustering of genes involved in reprogramming in GBM (TCGA cohort) based on high XBP1s or high RIDD activity (blue low levels, red high levels). **D)** mRNA expression of SOX2, POU3F2, OLIG2, SALL2 reprogramming TFs based on microarray fluorescence intensity in high XBP1s and high RIDD activity tumors (TCGA cohort). **E)** mRNA levels of SOX2 upon XBP1s silencing in primary and classical adherent GBM lines (RADH85/87 and U251 respectively). (ns): not significant; (*): p<0.05; (**): p<0.01; (***): p<0.001.

### XBP1s-regulated miR148a represses SOX2 and sustains GBM cell differentiation

We next sought to identify the mediating factor between XBP1s induction and reprogramming TFs downregulation and hypothesized that these factors might be miRNAs (Figure 6A, (18)). Using miRNA sequencing, we first identified miRNAs whose expression is dependent on the IRE1/XBP1s signaling axis and then investigated the expression of those miRNAs in tumors stratified based on high XBP1s or high RIDD activity (Figures 6B-C). This indicated that miR21 and miR148a were the two miRNAs presenting the highest expression in XBP1s high tumors. Interestingly, miR148a was identified previously as a transcriptional target of XBP1s using chromatin immunoprecipitation (19) and SOX2 mRNA presents miR148a binding sites **(Figures S6A-D)**. To further document the IRE1-dependent control of miR148a expression, we first showed that miR148a levels were decreased in IRE1 signaling deficient cells when compared to parental cells (Figure 6D). This also corresponded to conditions where the upregulation of SOX2 and other TFs was observed (Figures 1-3). To consolidate the link between the expression of miR148a and XBP1s, we next measured miR148a levels in cells silenced for XBP1s and we observed significant miR148a downregulation (Figure 6E). We next artificially upregulated the expression of miR148a in IRE1 signaling deficient cells using miR148a mimics and observed a downregulation of SOX2 as well as the majority of the other TFs as opposed to the upregulation we would normally observe (Figures 6F, **S6E**). Furthermore, to verify the link between XBP1s, miR148a and SOX2 signaling we overexpressed XBP1s in IRE1 deficient and parental cells. In these conditions we observed the subsequent upregulation of miR148a expression and the downregulation of SOX2 and other TFs in GBM cell lines (Figures 6G, **S6F**). A final validation of this signaling cascade was achieved by overexpressing XBP1s in IRE1 deficient cells and also treat the cells with miR148a inhibitors. As hypothesized, we observed an upregulation of XBP1s compared to untreated cells, a downregulation of miR148a and the subsequent upregulation of SOX2 (Figure 6H). Hence, the expression of reprogramming TFs including SOX2 is downregulated by miR148a, clarifying a relationship previously documented in the literature (20), and miR148a is directly induced by XBP1s (19). Therefore we delineate a novel IRE1-dependent signaling cascade that maintains GBM cell differentiation.

**Figure 6.**
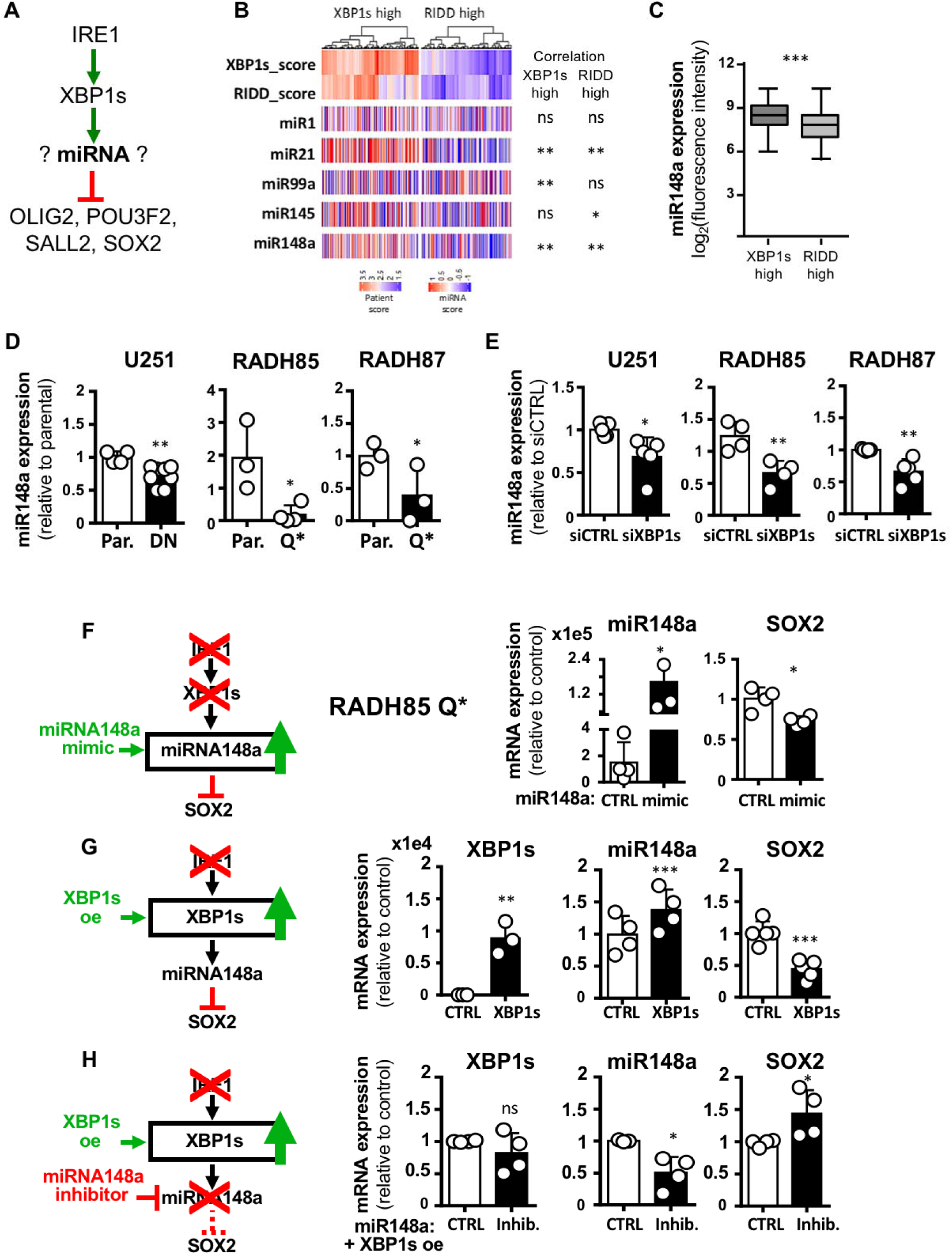
XBP1s-dependent expression of miR148a prevents GBM cell reprogramming. **A)** Schematic representation of hypothesis of effect of IRE1 signaling on reprogramming TFs. **B)** Hierarchical clustering of miRNAs in GBM (TCGA cohort) confronted to high XBP1s or high RIDD activity (blue low levels, red high levels) with the best 5 candidates shown. **C)** mRNA expression of miR148a based on microarray fluorescence intensity in high XBP1s and high RIDD activity tumors (TCGA cohort). **D)** miR148a expression in adherent lines U251, RADH85/87 expressing DN or Q* forms of IRE1 normalized to parental. **E)** miR148a expression in adherent lines U251, RADH85/87 transiently deficient for XBP1s through siRNA transfection compared to control. **F)** SOX2 and miR148a expression levels in RADH85 IRE1 Q* expressing cells in the presence of miR148a mimic compared to control. **G)** XBP1s, SOX2 and miR148a expression levels in RADH85 IRE1 Q* expressing cells, over-expressing XBP1s compared to control. **H)** XBP1s, SOX2 and miR148a expression levels in RADH85 IRE1 Q* expressing cells, over-expressing XBP1s, in the presence of miR148a inhibitors compared to control. (ns): not significant; (*): p<0.05; (**): p<0.01; (***): p<0.001.

### IRE1 signaling is essential for the maintenance of GBM differentiation *in vivo*

We next documented this hypothesis in a murine GBM model. To this end we measured the levels of XBP1s, miR148a and SOX2 in IRE1 knockout (KO) murine GL261 GBM cells. We found that the effect seen in human cells was recapitulated in murine GBM cells with miR148a downregulation and subsequent SOX2 upregulation following an expected XBP1s downregulation (Figures 7A, **S7**). Next, following intracranial injections of parental or IRE1-KO GL261 cells in the brains of C57BL/6 mice, tumors were allowed to grow. The brain was harvested and multiple sections of the tumors were stained for the stem marker MSI1 (Figure 7B) and quantified (Figure 7C). This system was also probed pharmacologically using a similar approach except that 14 days post-injection, tumors were surgically resected and a gel implant containing DMSO or MKC was placed in the tumor cavity (Figure 7D). Stem cell marker MSI1 staining was increased in both tumor and tumor periphery in both IRE1 KO (Figure 7B, C) or MKC (Figure 7E, F) conditions compared to control or DMSO conditions, respectively. No difference was observed in the opposite hemisphere parenchyma. We have thus demonstrated that IRE1 inhibition leads to an increase in the presence of the stem cell marker MSI1 in an *in vivo* model of GBM.

**Figure 7.**
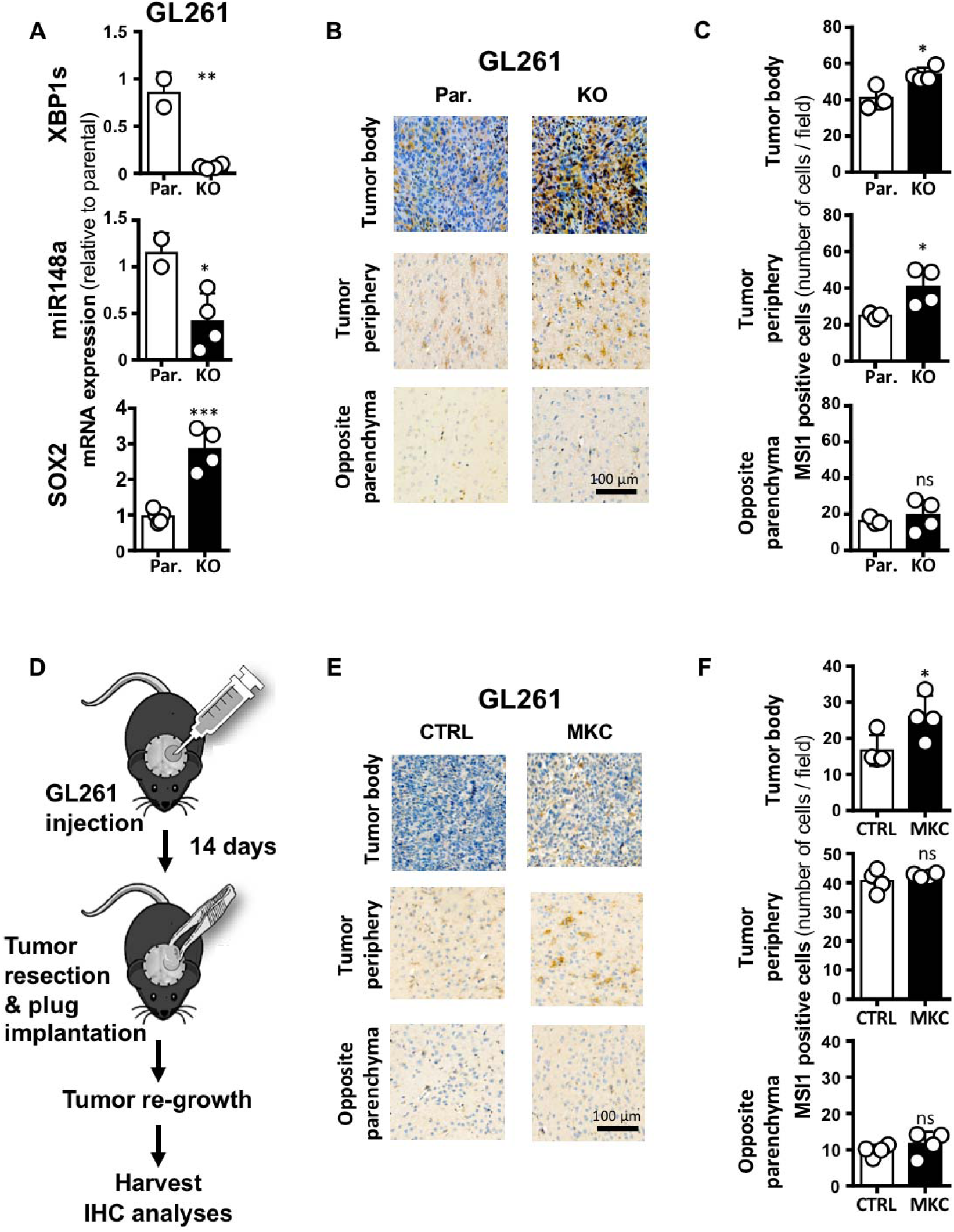
IRE1-dependent control of GBM stemness reprogramming *in vivo*. **A)** XBP1s, SOX2 and miR148a expression levels in GL261 IRE1 KO cells, compared to parental. **B)** Representative sections of tumors grown from GL261 parental or GL261 IRE1-KO cells injected in the brain of orthotopic syngeneic mouse model, stained for MSI. Sections from tumor body, tumor periphery and opposite to tumor brain parenchyma shown. **C)** Quantification of MSI1 positive cells (3-4 tumors/condition, 10 random fields/tumor/condition quantified, two independent counts). **D)** Orthotopic syngeneic mouse GBM model and peri-operative treatment with MKC (plug). **E)** MSI1 staining (stem marker) of tumor body or periphery and opposite to tumor site hemisphere parenchyma. This was performed in GL261 derived tumors treated with control or MKC plugs. **F)** Quantification of MSI1 positive cells (3 tumors/condition, 10 random fields/tumor/condition quantified, three independent counts). (ns): not significant; (*): p<0.05; (**): p<0.01; (***): p<0.001.

## Discussion

Our analysis of IRE1 signaling in GBM shows that the IRE1/XBP1s/miR148a signaling axis plays a major role in the maintenance of a differentiated cancer cell phenotype and paramount in absentia to the ability of cancer cells to revert to a stem-like phenotype. Using both GBM transcriptome data from the TCGA and animal models we show that reduced IRE1 activity in tumor cells correlated with enhanced expression of stem cells markers (Figures 1-3). Furthermore, using cultured cells, we demonstrate that using both genetic and pharmacological inhibition of IRE1 activity promotes differentiated cells to revert into more cancer stem-like cells (Figures 2, 3). We also delineate the molecular mechanism by which IRE1 exerts its differentiating functions. Indeed, we show that activation of the IRE1/XBP1s pathway controls the expression of miR148a, which in turn represses the expression of transcription factors involved in stemness (e.g. SOX2) (Figures 4, 5). Our focus on miR148a was motivated by the demonstration that this miRNA was identified as i) as a transcriptional target of XBP1s (19) and ii) a potential binder of SOX2 mRNA (**Figure S6**). Interestingly miR148a promotes plasma cell differentiation by targeting germinal center TFs MITF and BACH2 (21), which thus could also occur under the control of XBP1s, a major player in this process (22).

The role of the different branches of the UPR^ER^ has been shown in many instances to participate in the regulation of the stem-differentiated phenotype in various models. Indeed, pluripotent stem cells undergo various proteostatic stresses during reprogramming and as such the UPR^ER^ is of particular importance as its transient activation enhances these properties (23). In this model the important contribution of XBP1s was also observed (23). One might hypothesize that similar pathways could also contribute to the appearance of stem-like cells in diseases. Of particular clinical interest is the contribution of CSC to disease progression and relapse (24) and in this context, UPR^ER^ signaling was found to alter the CSC compartment in various solid cancers. Indeed, the constitutive activation of XBP1s in triple negative breast cancer (TNBC) was shown to increase the CSC compartment (6). In addition, in GBM, it was recently shown that the PERK arm of the UPR^ER^ was responsible for the attenuation of SOX2 expression upon acute ER stress and therefore facilitate CSC differentiation (25). The role of the UPR^ER^ on the stem-like vs. differentiated state of tumor cells was further illustrated through the involvement of the ATF6 arm which was shown to sustain dormant cells survival through specific pathways including RHEB and AKT (26). These data indicate that UPR^ER^ signaling might play a significant role in the balance between CSC and differentiated cancer cells, and exhibit different properties in various cancer types as well as various contributions of the different arms.

It is well established that heterogeneity is a major barrier to successful treatment in GBM. Moreover, de-differentiation of GBM cells to a stem-like phenotype compounds chemo-resistance. The contribution of IRE1 signaling in GBM development and outcome has been widely documented (4, 7, 8, 27). We have previously stratified GBM tumors according to IRE1 activity and XBP1s or RIDD activity (XBP1s conferring worse prognosis) (4). Here we provide evidence that IRE1 inhibition and particularly XBP1s inhibition might promote differentiated tumor cells to revert to a more stem-like phenotype, an effect that has been linked with chemo-resistance and oncogenesis in general in GBM (28). As such our results point to two distinct opportunities for utilizing IRE1 signaling as therapeutic target in GBM. Firstly, patients with XBP1s low expressing tumors could benefit from IRE1 inhibition as GBM cells would have little capacity to utilize this signaling pathway for reverting to a stem phenotype; compounded by a UPR disruption that would overload a stress response mechanism already having to deal with a hostile microenvironment and anti-cancer treatments. Indeed a recent study has shown that the IRE1-XBP1s axis is important in the adaptation to stress of leukaemic and healthy haematopoietic stem cells, as they enhance the ability of these cells to overcome ER stress and survive, promoting carcinogenesis (29). This information compounds the second and potentially more important outcome of our study that shows the attractiveness of IRE1 targeting therapeutics as adjuvant therapy alongside the currently established trident of surgery, chemotherapy and radiotherapy in GBM. The rationale would be that IRE1 targeting can sensitize GBM cells to therapy as it would weaken their responses and disallow them the time to adapt to the hostile conditions afforded by chemotherapy. As such patients would benefit by not only potential drops in the rates of tumor re-emergence but also of reduced need for repeated therapy doses, which by default carry unfavorable toxicity profiles.

In conclusion, having demonstrated the importance of IRE1 in GBM development, prognosis and aggressiveness; we described i) a direct link of IRE1 signaling to differentiated cells reprogramming to stem-like cells in cell culture and mouse models and ii) the correlation between low IRE1 activity and the enrichment in CSC markers in patients tumors. Thereafter we delved further into the mechanisms of this IRE1 control and propose a novel concept where inhibition of XBP1s signaling induces the expression of transcription factors involved in GBM reprogramming and showcase the downstream miR148a signaling that makes this possible. Our work not only adds to the repertoire of IRE1 activity in GBM but also offers scope for patients’ stratification and combination therapy development with IRE1 targeting at its epicenter. It achieves this whilst reinforcing the need for strict pharmacovigilance when considering the risks of novel therapeutic design and builds upon our previous work in the field of translational neuro-oncology and ER biology to provide an ever more detailed landscape of IRE1 involvement and thus scrutinize the exploitability of the modulation of its function.

## Materials & Methods

### Reagents and antibodies

All reagents not specified below were purchased from Sigma-Aldrich (St Quentin Fallavier, France). Recombinant human BMP4 protein was obtained from R&D Systems (Lille, France); MKC8866 from Fosun Orinove (Suzhou, China); siRNA targeting XBP1, miR148a inhibitors, mimics and controls (miRvana) were purchased from Thermo Fisher Scientific (Montigny le Bretonne, France). For flow cytometry, antibodies against human NG2, nestin, O4, SOX2, and TUB were obtained from R&D Systems, Biotechne (Lille, France); anti-A2B5 from Miltenyi Biotec (Paris, France); anti-GFAP from eBioscience (Thermo Fisher Scientific), anti-MSI1 from Chemicon (Merck Millipore, Molsheim, France) (**Table S1**).

### Cell culture and treatments

U87MG (ATCC) and U251MG (Sigma, St Louis, MO, USA) cells were authenticated as recommended by AACR (http://aacrjournals.org/content/cell-line-authentication-information) and tested for the absence of mycoplasma using MycoAlert® (Lonza, Basel, Switzerland) or MycoFluor (Invitrogen, Carlsbad, CA, USA). Primary GBM cell lines were obtained as described in (16). U87, U251 and primary RADH GBM cells were grown in DMEM Glutamax (Invitrogen, Carlsbad, CA, USA) supplemented with 10% FBS. Primary GBM stem-like cell lines (RNS) were grown in DMEM/Ham’s:F12 (Life Technologies) supplemented with B27 and N2 additives (Life Technologies), EGF (20 ng/ml) and basic FGF (20 ng/ml) (Peprotech, Neuilly-sur Seine, France). Primary RADH85, RADH87, RNS85 and RNS87 were stably transfected at MOI = 0.3 with pCDH-CMV-MCS-EF1-Puro-copGFP (System biosciences) empty vector (EV), pCDH-CMV-MCS-EF1-Puro-copGFP containing IRE1α wild-type sequence (WT), or mutated sequence (Q780*). These cells were selected using 2 μg/ml puromycin, and polyclonal populations were tested for GFP expression. Transfections of GBM primary cell lines with IRE1 WT and Q780* were performed using Lipofectamine LTX (Thermo Fisher Scientific), according to the manufacturer’s instructions. Murine GBM GL261 IRE1 KO was generated as described in (30) using specific sgRNAs to the mouse IRE1 (Fwd#1: 5’-CACCGCAGGGTCGAGACAAACAACA-3’ and Rev#1: 5’-AAACTGTTGTTTGTCTCGACCCTGC-3’; Fwd#2: 5’-CACCGCAAAATAGGTGGCATTCCAG-3’ and Rev#2: 5’-AAACCTGGAATGCCACCTATTTTGC-3’. GL261 cells were grown in DMEM Glutamax supplemented with 10% FBS. For XBP1s overexpression, cells were transiently transfected for 48 hours with XBP1s plasmid (Addgene, Teddington, United Kingdom) using Lipofectamine 2000 (Thermo Fisher Scientific), according to the manufacturer’s instructions. For siRNA XBP1s experiments, cells were transiently transfected for 48 hours with control and siRNA targeting XBP1s (Eurofins Genomics, Les Ulis, France, 5‘ AGAAGGCUCGAAUGAGUG 3’) using Lipofectamine iRNA max (Thermo Fisher Scientific), according to the manufacturer’s instructions. For miR148a treatment, cells were transiently transfected for 48 hours with miR148a mimics or inhibitors (miRvana) using Lipofectamine iRNA max (Thermo Fisher Scientific), according to the manufacturer’s instructions.

### Transcriptomic data from TCGA

The publicly available GBM dataset of The Cancer Genome Atlas (TCGA) (Consortium et al., 2007; consortium, 2008) was assessed from the NCBI website platform (https://gdc-portal.nci.nih.gov) and was analyzed using the BioInfominer (31) analysis pipeline (e-Nios, Greece, https://bioinfominer.com), which performs statistical, semantic network, topological analysis, exploiting various biological hierarchical vocabularies, with the aim to unearth, detect and rank significantly altered biological processes and the respective driver genes linking these processes. Genes were considered significantly differentially expressed if the p value was below 0.05. To analyze the miRNA database, hierarchical clustering algorithms and Pearson correlation analyses were carried out using R packages.

### Quantitative real-time PCR

Total RNA was prepared using the TRIzol reagent (Invitrogen, Carlsbad, CA, USA). Semi-quantitative analyses were carried out as previously described (7, 8). All RNAs were reverse-transcribed with Maxima Reverse Transcriptase (Thermo Scientific), according to manufacturer protocol. qPCR was performed with a StepOnePlus™ Real-Time PCR Systems from Applied Biosystems and the SYBR Green PCR Core reagents kit (Takara, Ozyme, Saint Cyr L’Ecole, France). All RNAs for the miRNA investigation were transcribed using miScript RT kits and subsequent qPCR performed using miScript Primer Assays and miScript SYBR kits (Qiagen, Courtaboeuf). Experiments were performed with at least triplicates for each data point. Each sample was normalized on the basis of its expression of the *actin* gene. For quantitative PCR, the primer pairs used are described in **Table S2**.

### Mouse Intracranial injections

Eight-week old male C57BL/6 mice were housed in an animal care unit authorized by the French Ministries of Agriculture and Research (Biosit, Rennes, France - Agreement No. B35-238-40). All animal procedures met the European Community Directive guidelines (DIR MESR 13480) and were approved by the local ethics committee No. 007. The protocol used was as previously described (7). Cell implantations were at 2 mm lateral to the bregma and 3 mm in depth using GL261 cells. Mice were daily clinically monitored and sacrificed at first clinical signs. In the experiments with MKC plug, fourteen days post-injection, the tumor formed was maximally removed without killing the animal and a plug infused with MKC or DMSO control was implanted in the resection cavity. At first clinical signs, mouse brains were collected, fixed in 4% formaldehyde solution and paraffin embedded for histological analysis using anti-vimentin antibody (Interchim, Montluçon, France) to visualize the tumor masses.

### Immunohistochemistry analyses

Mouse tumor tissues were fixed in 4% neutral buffered formalin, embedded in paraffin, cut into 5-μm thick sections and mounted on slides. Immunohistochemistry (IHC) was carried out on the H2P2 imaging platform. For cancer stem cell immunodetection, the sections were incubated 1 hour at room temperature with anti-MSI1 antibody (1:500 dilution; Merk Millipore). Immunostaining was carried out using the BenchMark XT (Ventana Medical Systems, Illkirch Graffenstaden, France) with the OmniMap kit (a “biotin-free” system using multimer technology, Roche, Boulogne Billancourt, France) and a Tris borate EDTA pH8 buffer for antigen retrieval. Stainings were analyzed with an Axioplan 2 epifluorescent microscope (Zeiss, Marly-le-Roi, France) equipped with a digital camera Axiocam (Zeiss), and were converted on to digital slides with the scanner Nanozoomer 2.0-RS (Hamamatsu, Meyer Instruments, Houston, United-States of America). ImmunoRatio, the publicly available web application (http://153.1.200.58:8080/immunoratio/) for automated image analysis, was used to determine the number of MSI1-stained cells.

### Flow cytometry analyses

Cells were washed with PBS 2% FBS and incubated with saturating concentrations of human immunoglobulins and fluorescent-labelled primary antibodies for 30 min at 4°C. Cells were then washed with PBS 2% FBS and analyzed by flow cytometry using a NovoCyte NovoSampler Pro (ACEA Biosciences, Ozyme). The population of interest was gated according to its FSC/SSC criteria. Data were analyzed with the NovoExpress software (ACEA Biosciences). For intracellular staining, cells were permeabilized using BD Cytofix/Cytoperm and BD Perm/Wash reagents (BD Biosciences, Le Pont de Claix, France) according to the manufacturer’s instructions. Protein expression levels were given by the ration of mean of fluorescence obtained with the antibody of interest divided by the mean of fluorescence obtained with the isotype control (rMFI).

### Statistical analyses

Data are presented as mean ± SD or SEM (as indicated). Statistical significance (P < 0.05 or less) was determined using a paired or unpaired t-test or ANOVA as appropriate and performed using GraphPad Prism software (GraphPad Software, San Diego, CA, USA).

## Acknowledgements

We thank the Biosit histopathology H2P2 platform (Université de Rennes 1, France) and Florence Jouan for immunohistochemical analyses of tumor xenografts; and the Biosit ARCHE animal facility (Université de Rennes 1) for animal housing. This work was funded by grants from Institut National du Cancer (INCa PLBIO), Fondation pour la Recherche Médicale (FRM, équipe labellisée 2018) to EC; EU H2020 MSCA ITN-675448 (TRAINERS) and MSCA RISE-734749 (INSPIRED) grants to AS, EC, EC; PHC Maimonide 2017-2018 to BT and EC; the David R. Bloom Center for Pharmacy, the Dr. Adolph and Klara Brettler Center for Research in Pharmacology, German Israeli Fund (grant no. I-1471-414.13/2018) to BT; the Ministry of Science & Technology, Israel, The Ministry of Europe and Foreign Affairs, France & the Ministry of Higher Education, Research and Innovation, France, to BT and EC; PROMISE, 12CHN 204 Bilateral Greece-China Research Program of the Hellenic General Secretariat of Research and Technology and the Chinese Ministry of Research and Technology sponsored by the Program “Competitiveness and Entrepreneurship,” Priority Health of the Peripheral Entrepreneurial Program of Attiki to AC. DD is a Marie Curie early stage researcher funded by EU H2020 MSCA ITN-675448 (TRAINERS). JO was funded by a post-doctoral fellowship from “Région Bretagne”. JS is funded by Hardiman Fellowship and Science Foundation Ireland (SFI) and the European Regional Development Fund (Grant Number 13/RC/2073). MMM was funded by the Irish Research Council and an ARED International PhD fellowship from “Région Bretagne”.

## Author contribution

**DD** – conceptualization, methodology, investigation; **MM, JO, RP, PJLR, AO, JS, AP, BT** – methodology, investigation; **KV** – data curation, formal analysis; **AC** – data curation, formal analysis funding acquisition; **JBP, QZ** – resources; **AS, BT** – resources, funding acquisition; **EC** – supervision, conceptualization, project administration, funding acquisition; writing; **TA** – supervision, conceptualization, methodology, investigation, project administration, writing (https://www.casrai.org/credit.html<u>#</u>)

