## Supplemental Material for "Constitutive UPR^ER^ activation sustains tumor cell differentiation"

**Supplemental Materials**

**Figure Legends**

**Figure S1. Exploratory Data Analysis (EDA) of IRE1 signature association with the mRNA expression level of differentiation- and stem-related genes in TCGA-GBM patient cohort**. Normalized Affymetrix U133 microarray data (527 cases) and scaled by gene log2CPM RNAseq data (156 cases) were exploited for gene expression profiling. **A)** Unsupervised hierarchical clustering analysis (pearson complete linkage on standardized microarray data) representing the association of IRE1_38 signature with the expression profile of differentiation and stem markers. The XBP1s component of IRE1 signature is represented by the red bar and the RIDD component by the blue one. **B)** Contingency tables of the distribution of IRE1 activity groups (groups in rows) by the distribution of differentiation- and stem-activity groups (groups in columns), respectively. Patient grouping was based on the quantile-oriented XBP1s score (IRE1_sign score) (1) and the differentiation/stem enrichment score (Diff/Stem_sign score). The latter denotes the ratio between the number of over- (or down-expressed) differentiation or stem markers to the total number of differentiation or stem markers were examined, respectively. Each differentiation or stem marker was appointed as over- or down-expressed when its gene expression in the sample was larger than the median gene expression in cohort. Mid- state represents equal distribution between over- and down-expressed markers. Chi-square (χ2) hypothesis tests of independence were performed to analyze the correlation between IRE1activity and differentiation and stem marker activity groups. The p-value of each statistical test is surrounded by a red box. **C)** Hierarchical clustering of TCGA-GBM patients (RNAseq data) based on their IRE1_sign score coupled with the differentiation/stem enrichment score. Samples were clustered using Euclidean distance metrics with Ward.D2 linkage. **D)** Hierarchical clustering of TCGA-GBM patients (RNAseq data) using the expression profile of IRE1 signature coupled with the expression levels of differentiation and stem markers. Complete linkage and Pearson correlation was used as distance measure. Two crosstabs are also used to aggregate and jointly display the distribution of IRE1activity groups with the distribution of differentiation- and stem-activity groups. Fisher's Exact Test was additionally performed because of the small number of cases. IRE1 high activity is statistically significant correlated with high differentiation activity and low stem activity and vice versa. p-value <0.0001 and p-value <0.001 is denoted as .000 and .001, respectively, by the SPSS statistical software. **E)** Contingency tables of the distribution of IRE1 activity groups (groups in rows) by the distribution of differentiation- and stem-activity groups (groups in columns), respectively; as described in (Figure S1B). **F)** mRNA expression levels of reprogramming TFs in U87 cells expressing a dominant negative form of IRE1 compared to parental lines.

**Figure S2. Genetic perturbation of IRE1 effect on GBM cell reprogramming in U87 cells. A)** Differentiated GBM cell line U87 was cultured in neurosphere medium and passaged every 14 days. If the number of cells was under the initial number of cells seeded (10^6^), the culture was stopped. **B)** Phenotypic characterization of U87 parental and overexpressing DN form of IRE1 when grown in neurosphere media. **C)** Percentage of living cells and **D)** cell number per passage in differentiated U87/251 and RADH85/87 GBM cells when grown in neurosphere media and carrying a WT, DN or Q* form of IRE1 compared to parental lines. **E)** Heat map representation of fold change of mRNA expression of genes involved in reprogramming, stemness and differentiation normalized to parental in U87 lines expressing DN form of IRE1 when grown in neurosphere media. **F)** Protein expression of reprogramming, stemness and differentiation markers in these lines compared to parental lines. **G)** Clonogenicity of differentiated lines expressing DN form of IRE1 compared to parental lines when grown in neurosphere media. (ns): not significant; (*): p<0.05; (**): p<0.01; (***): p<0.001.

**Figure S3.** **Pharmacological inhibition of IRE1 effect on GBM cell reprogramming in U87 cells. A)** Schematic representation of GBM cell working model of differentiated to stem cell phenotype culture in the presence of MKC. **B)** Differentiated GBM cell line U87 was cultured in neurosphere medium in the presence of MKC, and passaged every 14 days. If the number of cells was under the initial number of cells seeded (10^6^), the culture is stopped. **C)** Phenotypic characterization of parental adherent U87 lines through culture in neurosphere medium treated with MKC or DMSO. **D)** Percentage of living cells and **E)** cell number per passage in differentiated U87/251 and RADH85/87 GBM cells when grown in neurosphere media in the presence of MKC. **F)** Heat map representation of fold change of mRNA expression of genes involved in reprogramming, stemness and differentiation normalized to parental in U87 lines when grown in neurosphere media in the presence of MKC or DMSO. **G)** Protein expression of reprogramming, stemness and differentiation markers in these lines compared to parental lines. **H)** Quantification of clonogenicity of single cell parental U87 lines when seeded in serum-free medium in the presence of MKC or DMSO. (ns): not significant; (*): p<0.05; (**): p<0.01; (***): p<0.001.

**Figure S4.** **Role of IRE1 in GBM stem-to-differentiated state reprogramming proof of concept. A)** Schematic representation of GBM cell working model of stem-to-differentiated cell phenotype culture in the presence or absence of MKC. **B)** Phenotypic characterization of RNS85/87 parental and overexpressing WT or Q* forms of IRE1 when grown in FCS containing media (RNS + BMP4) with comparator panels when grown in neurosphere media (RNS). **C)** Protein expression of differentiation markers in RNS85/87 over 10 days in the presence of BMP4. **D)** Heat map representation of fold change of mRNA expression of genes in RNS85/87 in the presence of BMP4 for the final experimental conditions as dictated by (C). **E)** Protein expression of differentiation markers in RNS85/87 in the presence of BMP4 for the final experimental conditions as dictated by (C). (ns): not significant; (*): p<0.05; (***): p<0.001.

**Figure S5.** **XBP1s downregulation effects on reprogramming TFs in GBM cells.** SALL2, POU3F2, OLIG2 mRNA expression in parental adherent lines U251, RADH85/87 transiently deficient for XBP1s through siRNA transfection compared to control. (NA): no amplification; (ns): not significant; (*): p<0.05; (**): p<0.01.

**Figure S6.** **XBP1s downregulation and miR-148a overexpression have opposite effects on reprogramming TFs in GBM cells. A-D)** Sequence based evidence of potential SOX2/miR148a interaction. **A)** miR148a secondary structure. **B)** miR148a 5’ and 3’ strand sequences. **C)** SOX2 mRNA map with indications of sequence homology and hence binding site probability of both 3’ and 5’ strands of miR148a. Two sites are identified per strand, with miR148a 5’ potentially binding on both the 5’ and 3’ UTR of SOX2 mRNA whilst miR148a 3’ occupying two sites on the 3’ UTR. **D)** Species homology across human, monkey, rat and mouse of SOX2 and miR148a sequence complementarity of all four sites identified in C). **E)** OLIG2, POU3F2, SALL2 and SOX2 mRNA expression levels in U251, RADH85/87 IRE1 DN and Q* expressing cells in the presence of miR148a mimic compared to control. **F)** SOX2, SALL2, POU3F2 and OLIG2 mRNA expression levels in U87/251 and RADH85/87 IRE1 DN/Q* expressing cells, over-expressing XBP1s compared to control. (NA): no amplification; (ns): not significant; (*): p<0.05; (**): p<0.01; (***): p<0.001.

**Figure S7. XBP1s controls expression levels of TFs involved in reprogramming in murine GBM GL261 cell line.** OLIG2, POU3F2, SALL2 mRNA expression levels in GL261 cells, null for IRE1 (KO) compared to parental. (NA): no amplification; (ns): not significant; (***): p<0.001.

**Table S1:** Antibodies used in flow cytometry and immunohistochemistry experiments

| **Antigen** | **Usage in this study*** | **Fluorescence / dilution** | **Clone** | **Species / isotype** | **Company** | **Reference** |
| --- | --- | --- | --- | --- | --- | --- |
| A2B5 | FC | APC | 105HB29 | mouse IgM | Miltenyi Biotec | 130-093-562 |
| GFAP | FC | AF488 | GA5 | mouse IgG1 | eBioscience | 53-9892-82 |
| MSI1 | IHC | 1/500 | - | rabbit pAb | Chemicon | AB5977 |
| Nestin | FC | FITC | #96908 | mouse IgG1 | R&D Systems | IC1259F |
| NG2 | FC | FITC | LHM-2 | mouse IgG1 | R&D Systems | FAB2585F |
| O4 | FC | AF594 | O4 | mouse IgM | R&D Systems | FAB1326T |
| SOX2 | FC | AF594 | #245610 | mouse IgG2a | R&D Systems | IC2018T |
| TUJ1 | FC | APC | #TuJ-1 | mouse IgG2a | R&D Systems | IC1195A |

* FC for flow cytometry; IHC for immunohistochemistry.

**Table S2:** Primers sequences used in the Q-PCR experiments

| **Target gene** | **Sense*** | **Primer sequences** |
| --- | --- | --- |
| actin | Fwd | 5′-AGAGCTACGAGCTGCCTGAC-3’ |
|  | Rev | 5′-AGCACTGTGTTGGCGTACAG-3’ |
| BMI1 | Fwd | 5′-CACCAGAGAGATGGACTGACAA-3’ |
|  | Rev | 5′-AGGAAACTGTGGATGAGGAGAC-3’ |
| CD44 | Fwd | 5′-TGACACTGTCCAAAGGTTTTC-3′ |
|  | Rev | 5′-TCACTAATAGGGCCAGCCTC-3′ |
| CD133 | Fwd | 5′-TTTTGGATTCATATGCCTTCTGT-3’ |
|  | Rev | 5′-ACCCATTGGCATTCTCTTTG-3′ |
| GFAP | Fwd | 5′-GGAAGCGAACCTTCTCGATGTA-3’ |
|  | Rev | 5′-TAGGTGGCGATCTCGATGTCC-3’ |
| MBP | Fwd | 5′-AGGTCTCGTTCCGTGCTG-3’ |
|  | Rev | 5′-GCCACCATCCCTTGTGAG-3’ |
| NANOG | Fwd | 5′-GAAATACCTCAGCCTCCAGC-3′ |
|  | Rev | 5′-GCGTCACACCATTGCTATTC-3’ |
| NESTIN | Fwd | 5′- GCAGCAGGAAATATGGGAAG-3′ |
|  | Rev | 5′-TCTCATGGCTCTGGTTTTCC-3’ |
| NG2 | Fwd | 5′-CACACAGAGGAACCCTCGAT-3’ |
|  | Rev | 5′-CTTCAGCGAGAGAGGAGCACTT-3’ |
| OCT4 | Fwd | 5′-TCTCCCATGCATTCAAACTGAG-3′ |
|  | Rev | 5′-CCTTTGTGTTCCCAATTCCTTC-3′ |
| OLIG2 | Fwd | 5’-GGTAAGTGCGCAATGCTAAGCTGT-3’ |
|  | Rev | 5’-TACAAAGCCCAGTTTGCAACGCAG-3’ |
| POU3F2 | Fwd | 5′-ACACTGACCGATCTCCACGCAGTA-3’ |
|  | Rev | 5′-GAGGGTGTGGGACCCTAAATATGAC-3’ |
| SALL2 | Fwd | 5′-CGGATACCCATTGTGTCCT-3’ |
|  | Rev | 5′-CAGCATTTGGCACAGACTTG-3’ |
| SOX2 | Fwd | 5′- GAGCTTTGCAGGAAGTTTGC -3′ |
|  | Rev | 5′- GCAAGAAGCCTCTCCTTGAA -3′ |
| TUB | Fwd | 5′-ACGACGCTGAAGGTGTTCAT-3’ |
|  | Rev | 5′-AGTGTGAAAACTGCGACTGC-3’ |
| XBP1s | Fwd | 5′-TGCTGAGTCCGCAGCAGGTG-3’ |
|  | Rev | 5′-GCTGGCAGGCTCTGGGGAAG-3’ |
| YKL40 | Fwd | 5′- CCTGCTCAGCGCAGCACTGT -3′ |
|  | Rev | 5′- GCTTTTGACGCTTTCCTGGTC -3′ |

* Fwd for forward; Rev for reverse.

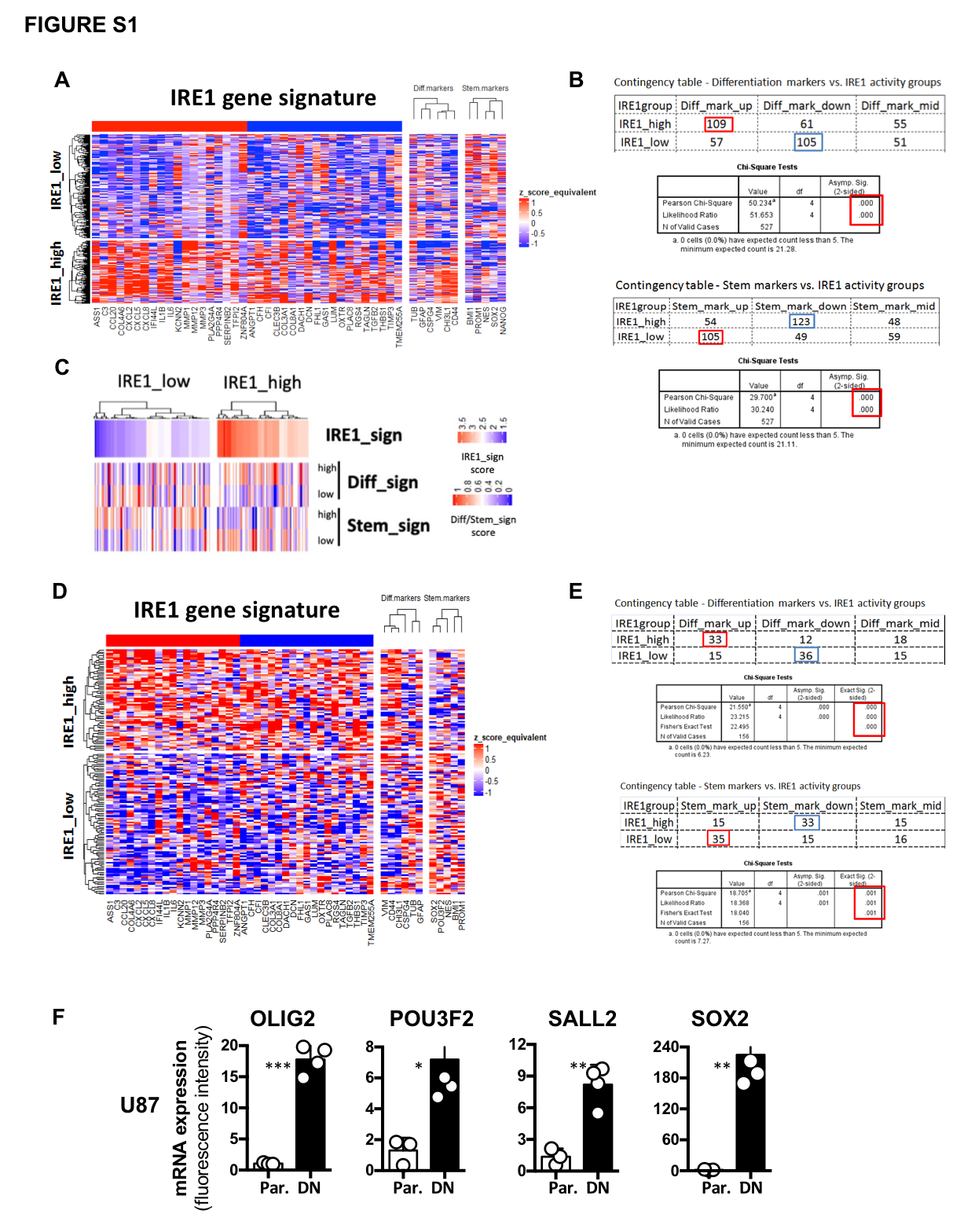

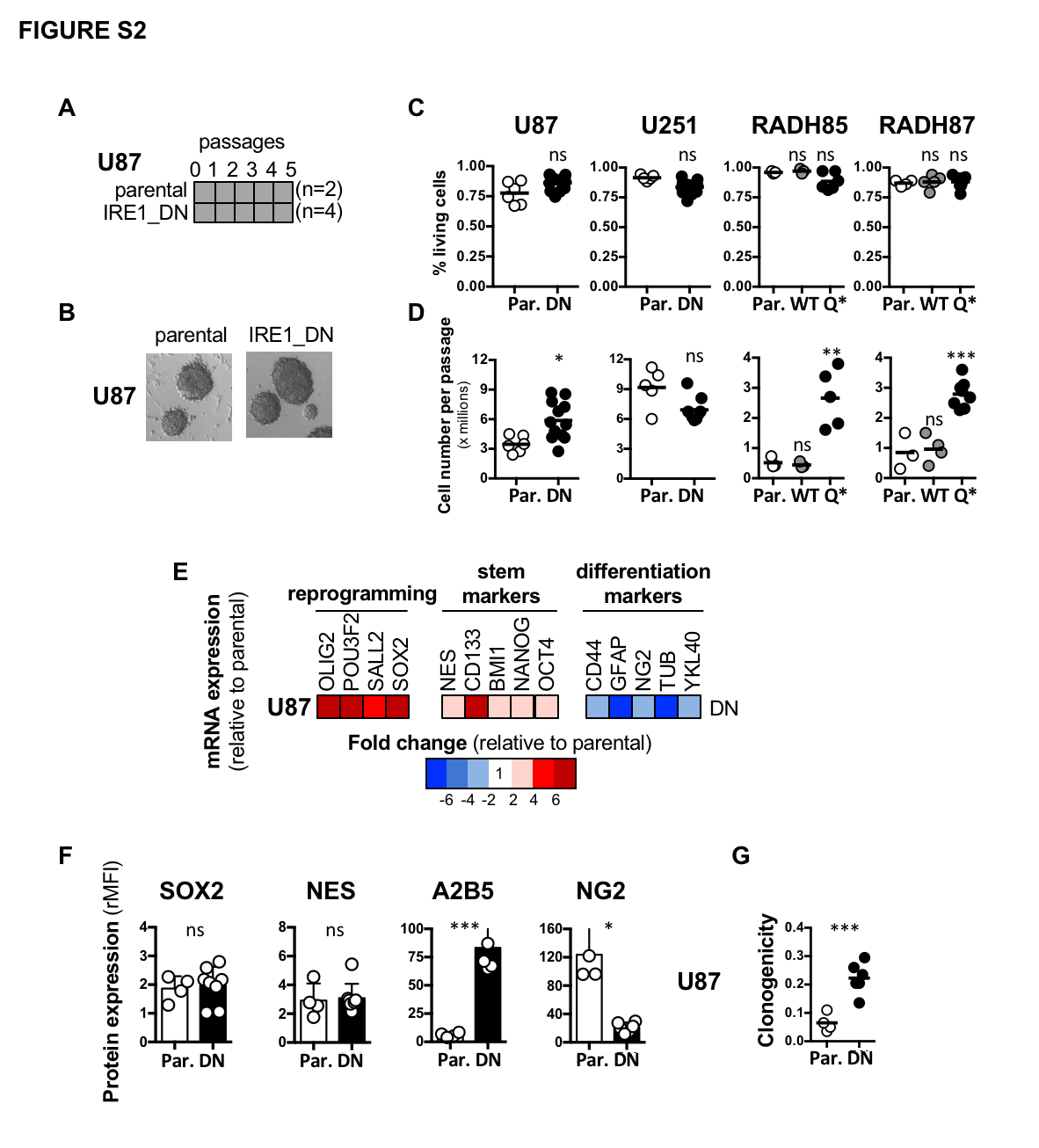

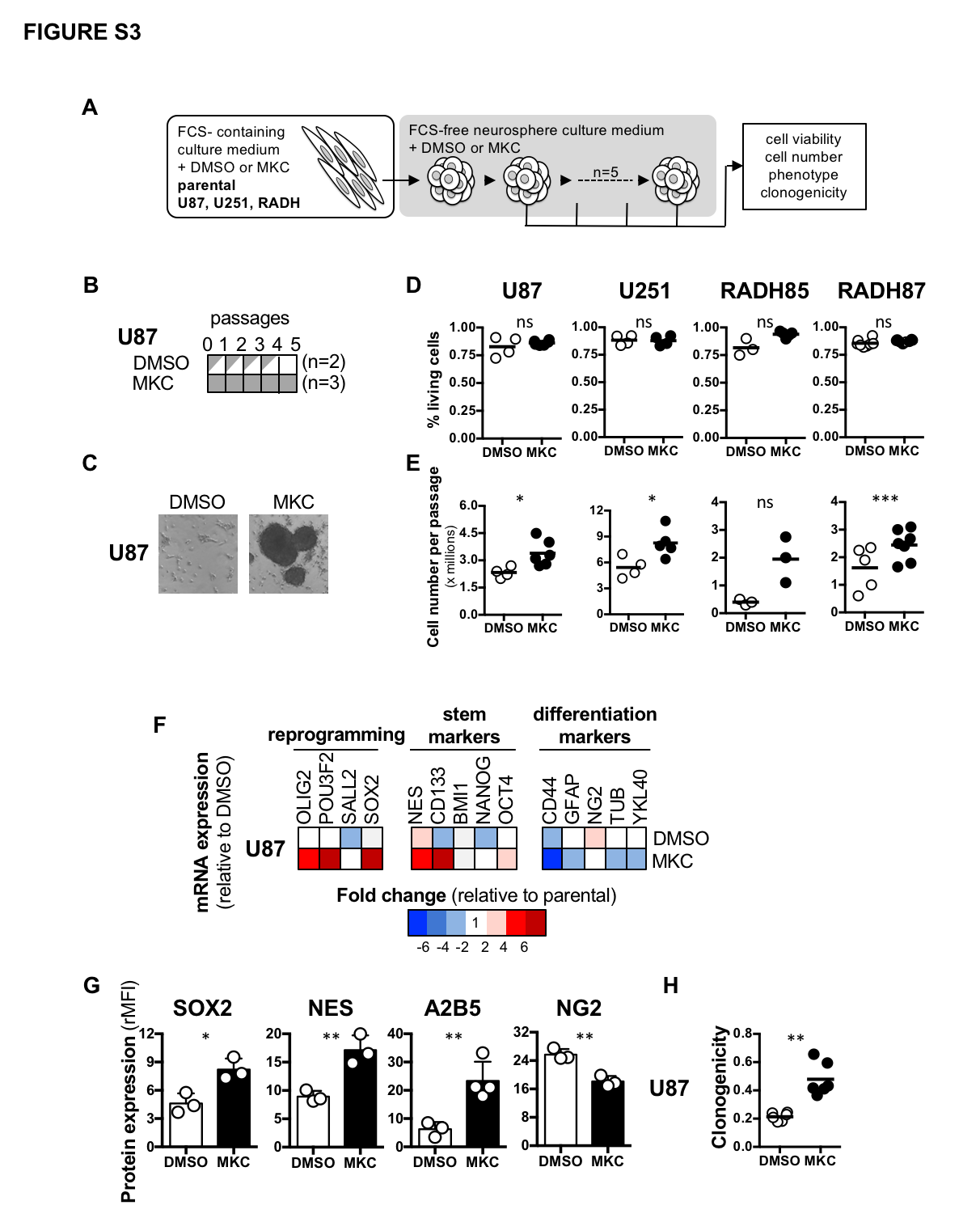

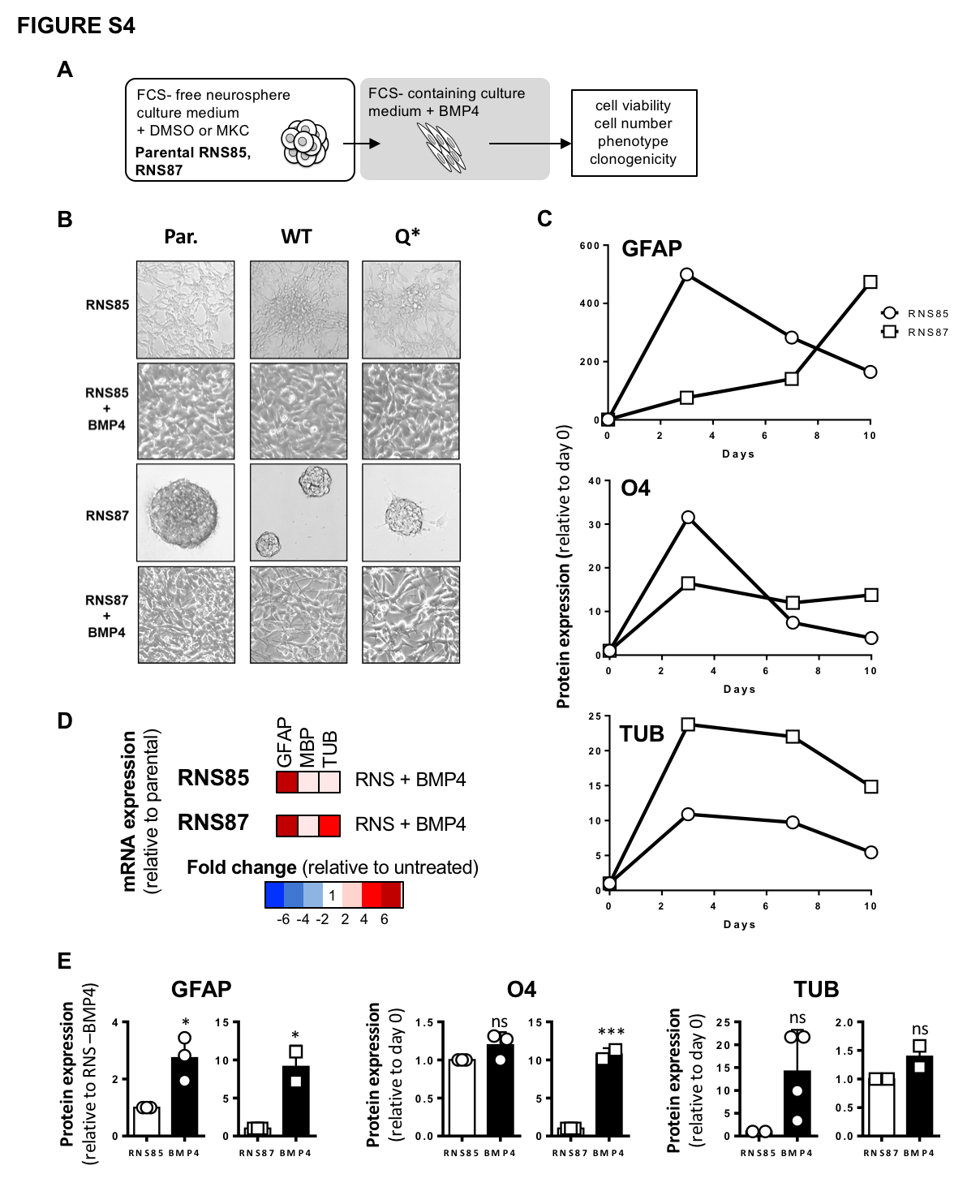

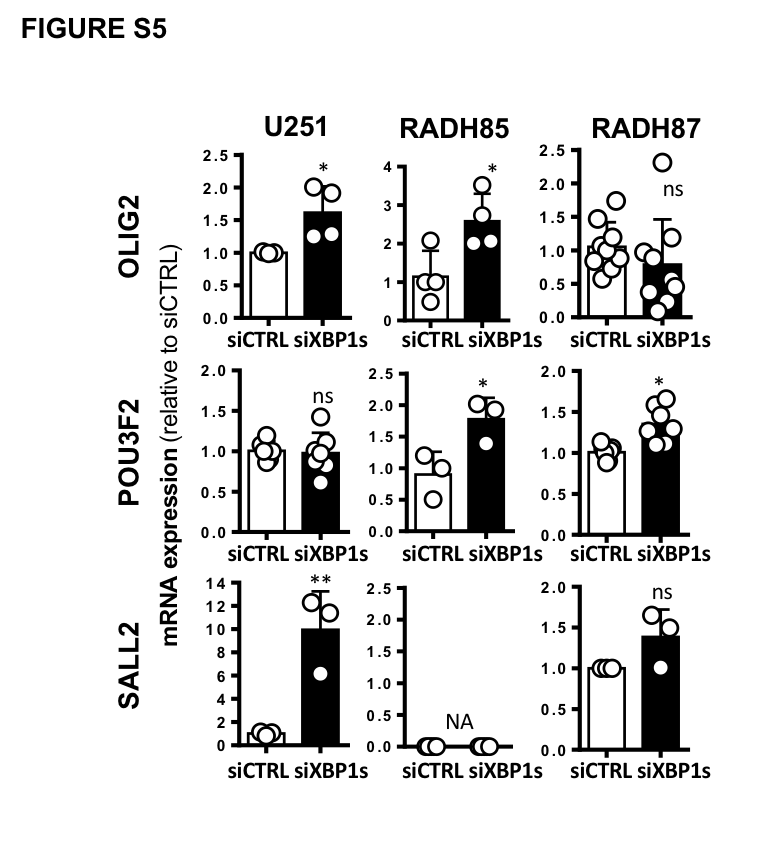

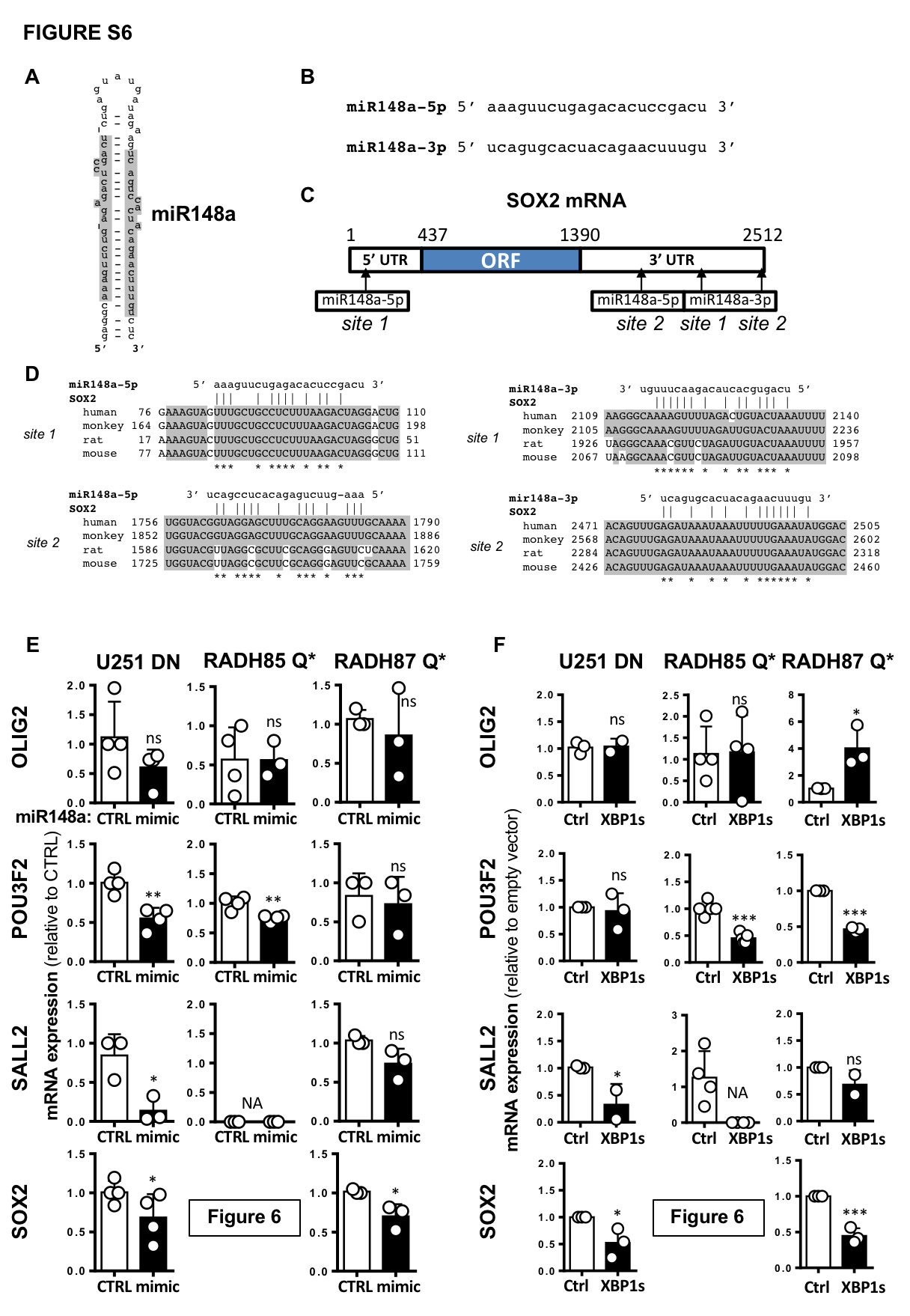

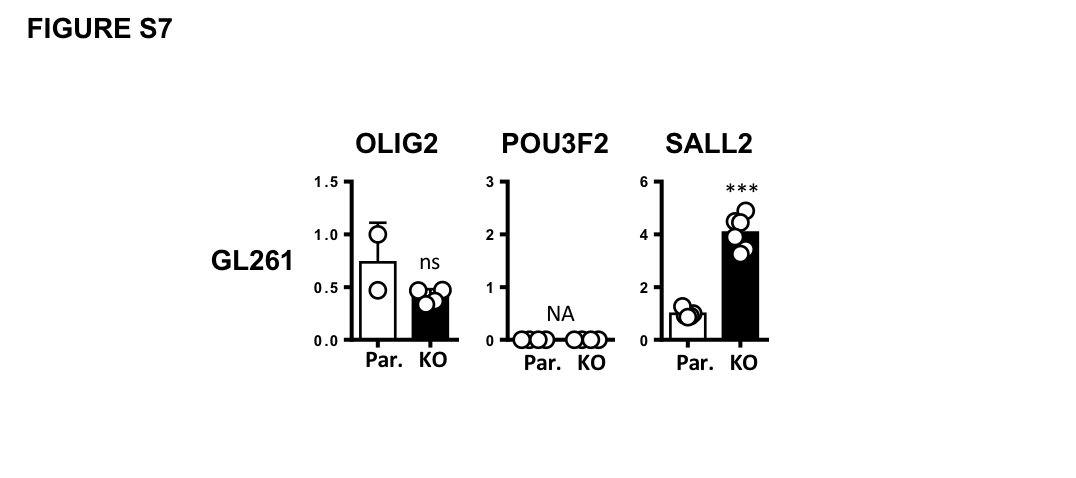
